# Substrate specificity and functional conservation of SWEET10 transporter in pineapple

**DOI:** 10.1101/2023.04.06.535793

**Authors:** Beenish Fakher, M. Arif Ashraf, Lulu Wang, Xiaomei Wang, Ping Zheng, Mohammad Aslam, Yuan Qin

## Abstract

- SWEET transporters are a unique class of sugar transporters that play vital roles in various developmental and physiological processes in plants.
- While the functions of SWEETs have been well established in model plants such as *Arabidopsis*, their functions in economically important fruit crops like pineapple have not been well studied.
- This study investigated the substrate specificity of pineapple SWEETs by comparing the protein sequences of known glucose and sucrose transporters in *Arabidopsis* to those in pineapple.
- Genome-wide approach and 3D structure comparison showed that the *Arabidopsis* SWEET8 homolog in pineapple, AcSWEET10, shares similar sequences and protein properties responsible for glucose transport. To determine the functional conservation of AcSWEET10, its ability to complement glucose transport mutants in yeast, its expression in stamens, and its impact on the microspore phenotype and seed set in transgenic *Arabidopsis* were analyzed.
- AcSWEET10 was found to be functionally equivalent to AtSWEET8 and plays a critical role in regulating microspore formation through the regulation of CalS5.
- Manipulating SWEET transporter activity could have important implications for improving fruit crop yield and quality.

## Introduction

Sugar is critical for the lives of all living organisms as it provides energy and carbon atoms for metabolic processes. Plants produce their sugar through photosynthesis, but the distribution of sugar from source to sink, sink to cell, and cell to cell requires specific sugar transporters (Hedrich *et al*., 2015). Multiple sugar transporters control the mobility and availability of sugar content in plants during their growth and development. To date, several families of sugar transporters have been discovered in plants, including Sucrose Uptake Transporters (SUTs/SUCs), Sugar Transport Proteins (STP) and Sugars Will Eventually be Exported Transporters (SWEETs) (Büttner & Sauer, 2000; Johnson & Thomas, 2007). SWEETs are the most recently discovered and functionally diversified sugar transporters. At the same time, SWEETs are evolutionary conserved based on the available sequenced plant genomes (Li *et al*., 2018; Xue *et al*., 2022). These transporters act as uniporters and are relatively pH-independent *in-vivo* conditions (Chen, L-Q *et al*., 2010; Chen *et al*., 2012). Interestingly, their subcellular localizations have been observed in the plasma membrane (Chen, L-Q *et al*., 2010; Chen, L-Q *et al*., 2015; Zhang *et al*., 2021), vacuolar membrane (tonoplast) and Golgi membranes (Chardon *et al*., 2013; Klemens *et al*., 2013; Chen, L-Q *et al*., 2015). Such diversification of SWEET transporter localization in eukaryotes or plants differs from prokaryotes. As prokaryotes lack membrane-bound organelles, the diversification of SWEET transporters’ localization provides a prime example of eukaryotic evolution. Additionally, the evolution of prokaryotic SWEETs (semiSWEETs) into eukaryotic SWEETs is thought to have arisen through gene duplication (Keller *et al*., 2014).

To date, significant progress has been made in understanding the transport route and mechanism of SWEET transporters through the use of crystal structures of semiSWEETs which served as a valuable foundation for the structural studies of SWEETs in eukaryotes (Xuan *et al*., 2013; Xu *et al*., 2014; Lee *et al*., 2015). Structurally, eukaryotic SWEETs are composed of two Triple Helix Bundle (THB) domains (TM1-TM2-TM3 and TM5-TM6-TM7), each of which is derived from semiSWEETs and separated by a linker helix (TM4). The N-terminal and C-terminal THBs share a similar sequence and are arranged in a parallel orientation (Chen, L-Q *et al*., 2010; Xuan *et al*., 2013). The distinct conformations of the structure allow them to be uniporter to access substrate (sugar molecules) from either the extracellular or cytosolic environments (Chen, L-Q *et al*., 2010; Guo *et al*., 2014; Feng & Frommer, 2015). The substrate-binding pocket is located above the center of the transmembrane region of the transporter proteins. Computational analysis has indicated that disaccharides such as sucrose, are able to easily pass through larger pockets. At the same time, monosaccharides such as glucose, due to their smaller size, can escape even if the pocket is partially open. However, extracellular residues of the transporter play a significant role in substrate recognition (Wang *et al*., 2014; Selvam *et al*., 2019).

The substrate preference of SWEET transporters is well explained through their evolution. Focusing on the evolution of SWEET transporters, the plant genome is known to possess approximately 20 paralogs of the SWEET gene family, which have been traditionally classified into four distinct clades, namely Clades I, II, III, and IV. Interestingly, the preference for sucrose and hexoses of SWEETs appear to be correlated with phylogeny; for example, SWEETs from clade III mediate sucrose transport. Functional characterization of SWEET transporters in the model plant *Arabidopsis* demonstrated that SWEETs are integral to seed filling (Chen, L-Q *et al*., 2015), nectar production (Lin *et al*., 2014) and pollen nutrition (Guan *et al*., 2008; Wang, Jiang *et al*., 2022).

Although, the fundamental ideas about the function, localization, and substrate specificity of SWEET transporters have been studied in the model plant *Arabidopsis thaliana*, the vastly diverse SWEET transporters available in the plant kingdom are entirely unexplained. In this study, we aim to expand the knowledge base of SWEET proteins by exploring the available information on SWEETs in pineapple, an economically important plant for which there is currently limited information. Instead of linear protein sequences, we utilized the transmembrane topology and AlphaFold to decipher the structural and physiological role of SWEETs in pineapple. This study combined the three-dimensional structure based transporter screening for their substrate specificity from the evolutionary perspective.

## Materials and methods

### Bioinformatic analysis

The protein sequences of pineapple were obtained from the Phytozome (https://phytozome.jgi.doe.gov/pz/portal.html). Protein sequences of *Arabidopsis*, rice *(Oryza sativa*), and maize were downloaded from TAIR (http://www.arabidopsis.org), the rice data center of China (http://www.ricedata.cn/gene/index.htm), and MaizeGDB (https://www.maizegdb.org/), respectively. The ExPASy (http://ca.expasy.org/prosite/) proteomics server was used for the physicochemical properties. WoLFPSORT (http://wolfpsort.hgc.jp) algorithm was utilized to predict the subcellular localization and TMHMM Server v.2.0 (http://www.cbs.dtu.dk/services/TMHMM/) was used to predict transmembrane helical domains available. The protein sequences were aligned, and the phylogenetic tree was constructed using MEGA (version 11.0) with the neighbor-joining (NJ) method (Tamura *et al*., 2013). The NJ tree was built with the ‘pairwise deletion’ option and the ‘Poisson correction’ model, and the internal branch reliability was assessed with a bootstrap test with 1000 iterations (Kumar *et al*., 2016). The transmembrane topologies were predicted and generated with Protter (Omasits *et al*., 2014).

### Homology-based modeling

The structure models of pineapple SWEET proteins in inward-facing states were built from Alpha fold (Jumper *et al*., 2021) and were visualized and analyzed using the CLC genomics workbench (version 22.0.2). The models were compared by using available structural templates: AtSWEET8 and AtSWEET13.

### Plant materials

*Arabidopsis* mutant (*Atsweet8*) was grown in a growth chamber in potted soil under a 16-h light/8-h dark regime at 22±2 °C. Transformation of *Arabidopsis* plants was done with *Agrobacterium* strain GV3101 using the floral dip method (Clough & Bent, 1998). Pineapple plants (MD2 variety) were acclimated in soil mix [peat moss: perlite = 2:1 (v/v)] in plastic pots in a walk-in growth chamber. The growth chamber was maintained at 25±2 °C with 16 h light/8 h dark photoperiod and 70% humidity, as reported earlier (Aslam *et al*., 2019). The pineapple anthers were collected and photographed at six developmental stages Wang *et al*. (2020)Wang *et al*. (2020)Wang *et al*. (2020)Wang *et al*. (2020), quickly frozen in liquid nitrogen, and stored at -80°C until RNA extraction.

### Total RNA isolation and quantitative-real time PCR analysis

Total RNA was isolated using the RNeasy kit (Qiagen, MD, USA), followed by treatment with DNaseI (Thermo Fisher Scientific, CA, USA) and reverse-transcribed using the ThermoScript RT-PCR kit (Thermo Fisher Scientific, CA, USA). The qRT-PCR reactions were set up with FastStart DNA Master SYBR Green I master mix (Takara, Japan). For each analysis, two technical replicates from three biological replicates were taken, and pineapple and *Arabidopsis EF1α* genes were used to normalize the mRNA levels. Finally, the fold change of genes was calculated using the 2^−ΔΔCT^ method (Livak & Schmittgen, 2001). Primers used in this study are listed in supplementary Table S3.

### RNA-Seq analysis

Transcriptome data of pineapple stamen developmental stages were used to investigate the expression level of SWEET genes (Wang *et al*., 2020). Briefly, the sequencing reads were aligned to the pineapple genome using TopHat v2.1.1 with default parameters. The transcript abundance was calculated as fragments per kilobase of exon model per million mapped reads (FPKM). The heatmap was generated using pheatmap R software based on log2 (FPKM+0.01).

### Transient expression of pineapple SWEETs in tobacco epidermal cells

The agrobacteria cultures of *35S::AcSWEET6:GFP*, *35S::AcSWEET8:GFP*, *35S::AcSWEET10:GFP*, *35S::GFP* (empty vector) and *35S::AtSWEET8:GFP* (as positive control) fusion constructs were pelleted and resuspended in the infiltration media (Bashandy *et al*., 2015). Using a needleless syringe, the suspension (OD 0.5) was infiltered into the abaxial surface of fully expanded tobacco leaves of four-week-old plants. GFP signals were detected at different time intervals between 48-96 h post infiltration (hpi) with the TCS SP8 microscope (Leica).

### Pollen viability analysis

Anthers were dissected from flower buds at stage 12 (before anthesis) and fixed in Carnoy fixative (absolute ethanol: chloroform: acetic acid, 6:3:1) for at least 2 h. Afterwards, anthers were stained in alexander stain (Ross *et al*., 2010). For aniline staining, anthers at the tetrad stage were gently fixed overnight in FAA. The anthers were then stained in 0.1 % (w/v) aniline blue in 0.1 M sodium phosphate (pH 9.0) and incubated in the dark for 1-2 h. Stained anthers were mounted on 30 % glycerol and viewed with a UV epifluorescence (365 nm excitation and a 420 nm long pass emission filter).

### Functional complementation of AcSWEETs in *Arabidopsis* (*sweet8*) mutant

PCR fragments of the complete *AcSWEET6*, *AcSWEET8*, and *AcSWEET10* coding sequences were amplified from flower-specific cDNA using specific primers. The fragments were cloned into the entry vector pENTR^TM^/D-TOPO® and then subcloned into the Gateway destination vector pGWB505 (Karimi *et al*., 2002) using LR Clonase II (Invitrogen). After confirmation through sequencing, the constructs were transformed into *Agrobacterium* and finally into homozygous *Atsweet8/rpg1 Arabidopsis* plants using the floral dip method. The transgenic plants were selected on media plates containing 50 mg l^−1^ hygromycin.

### Glucose transport assay in yeast

For the yeast transport assays, the *Saccharomyces cerevisiae* strain EBY.VW4000 (Wieczorke *et al*., 1999) was used as described previously (Fakher *et al*., 2022). The ORFs of *AcSWEET6*, *AcSWEET8*, and *AcSWEET10* were amplified using specific primer combinations. The amplified fragments were then cloned into the yeast expression vector, pGBKT7, and sequenced. The expression clones and an empty vector (without insert) were transformed into the yeast strain EBY.VW4000. Yeast transformants were then selected on a synthetic deficient medium without tryptophan (SD/-Trp) supplemented with 2 % (w/v) maltose as a carbon source. Transformed yeast cells were grown in SD/-Trp liquid medium with 2 % (w/v) maltose and were incubated overnight at 30°C until t he optical density at 600 nm (OD_600_) reached 0.5. The adjusted OD_600_ (∼0.2) with water was serially diluted (×10, ×100 and ×1000) and spotted onto SD/-Trp solid medium with 2% (w/v) maltose (control) and 2% (w/v) glucose. All the transformants were incubated at 30°C, and the growth was documented after 3 days in maltose media and 5-6 days in glucose media.

## Results

### 3D structure-based similarity among *Arabidopsis* and pineapple SWEET transporters

The degree of sequence identity among SWEETs can often correlate to their functional similarity. For example, maize ZmSWEET4c and its ortholog OsSWEET4 in rice are hexose transporters that play a role in seed filling and size determination (Sosso *et al*., 2015). Therefore, we first studied the close relatedness of SWEET transporters by constructing a phylogenetic tree using existing information from rice, grape, maize, *Arabidopsis*, and pineapple. Most pineapple SWEETs were closely linked to corresponding SWEETs from *Arabidopsis,* forming clades I, II, III, and IV (Figure 1). The average sequence identity of pineapple SWEETs was found to be 50%. To determine their potential target sugar for transport, the pineapple SWEETs were compared to well-characterized glucose (OsSWEET2b, AtSWEET8) and sucrose (AtSWEET13) transporters. The results revealed sequence similarities between pineapple SWEETs and known glucose and sucrose transporters, with AcSWEET1, 2, and 3 having a high sequence similarity to OsSWEET2b (Clade I). At the same time, AcSWEET5, 6, 7, 8, 9, and 10 showed similarities to AtSWEET8 (Clade II). AcSWEET11, 12, 13, 14, and 15 shared similarities with AtSWEET13 (Clade III), while AcSWEET16 and 17 had similarities to AtSWEET13 (Clade III). AcSWEET4 and 18 displayed lesser sequence similarities with AtSWEET8 and AtSWEET13 sequences (Figure 1, 2a and Table S1).

**Figure 1:**
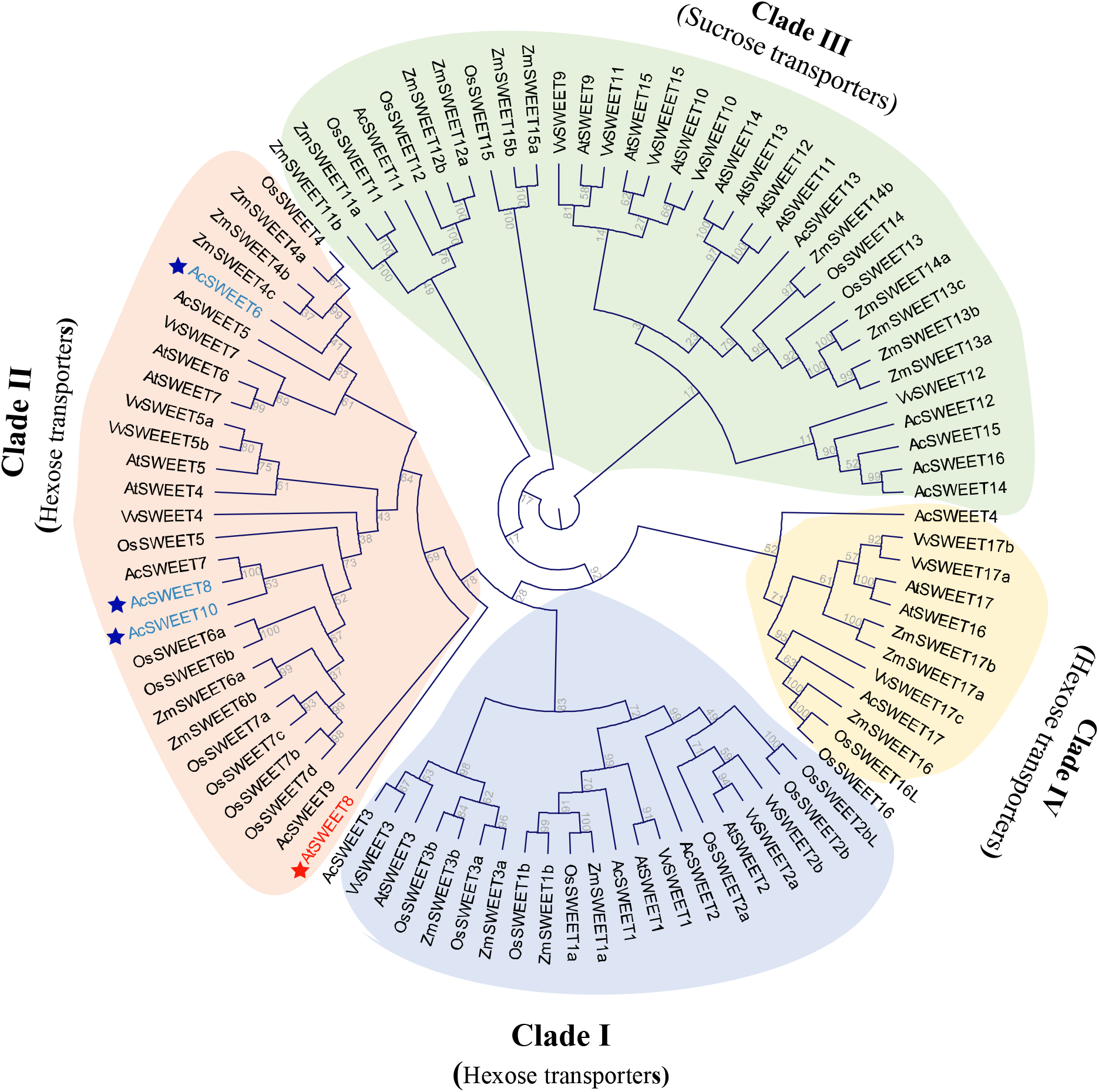
Phylogenetic tree of SWEET proteins from Pineapple (Ac), *Arabidopsis* (At), rice (Os), grape (Vv) and maize (Zm). The distances in the tree were calculated from multiple sequence alignment (CLC software) using the neighbor-joining method. Bootstrap values are displayed in percentages of 100. *Arabidopsis* SWEET8 protein is highlighted with red color and pineapple SWEET6/8/10 are represented with blue color.

**Figure 2:**
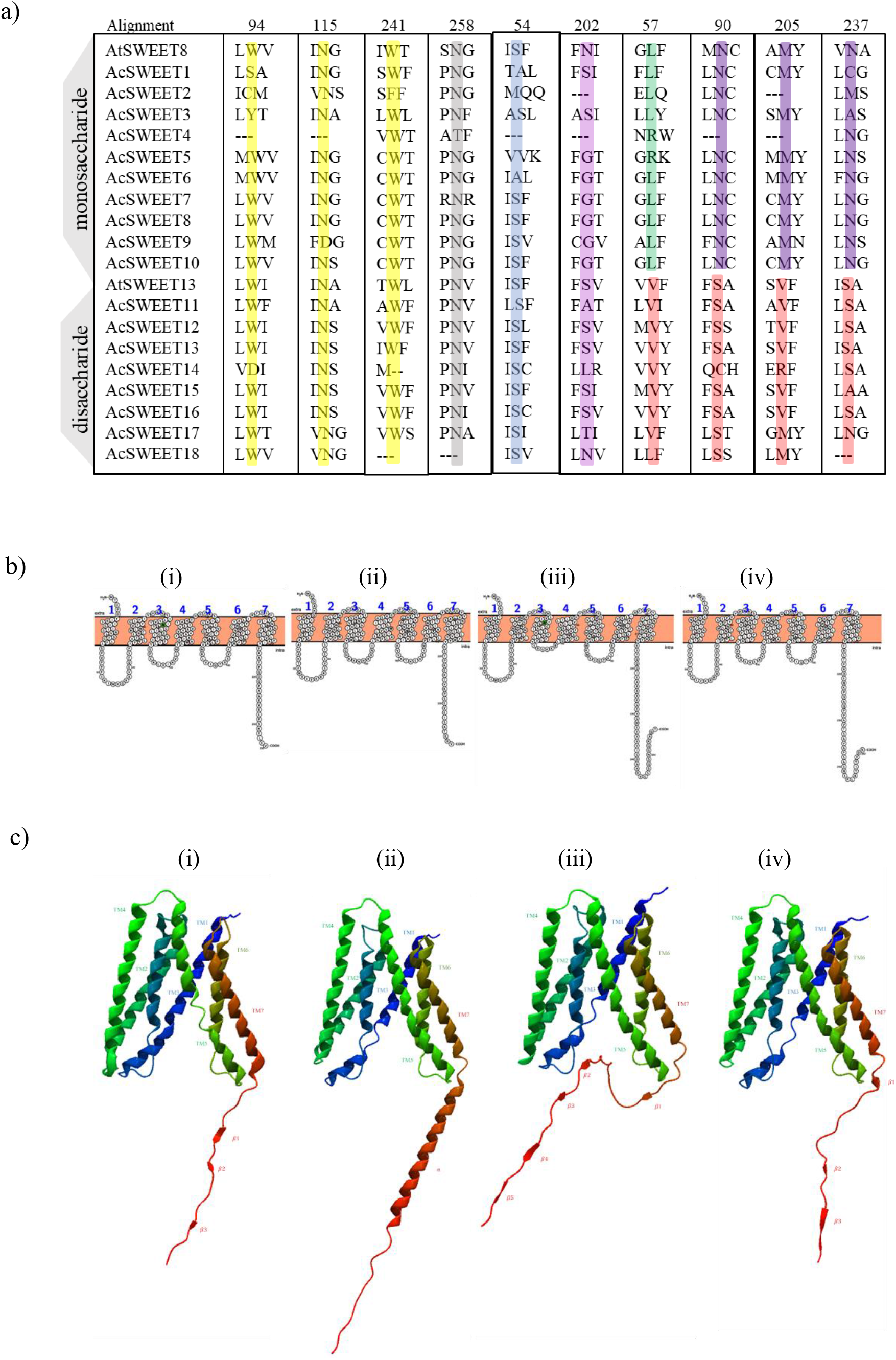
(a) Sequence alignment of pineapple SWEET proteins representing the four residues conserved among di- and monosaccharide-specific SWEETs are highlighted in cyan and green, respectively, numbers on top of the table represents AtSWEET13 (b) Predicted transmembrane topologies of selected SWEET proteins (c) Three-dimensional structures of SWEET proteins. In the figure, the numbers i, ii, iii and iv represent AtSWEET8, AcSWEET6, AcSWEET8, and AcSWEET10 proteins, respectively. The numbers on transmembrane helices (TM) are represented as TM1 to TM7. TM4 (linker helix) and the C-terminal tails are represented in blue and red colors respectively.

We then chose three pineapple SWEETs (AcSWEET 6, 8, and 10) from clade II for structural and functional characterization because they were similar to the previously well-characterized AtSWEET8 in terms of their physiochemical traits and projected sub-cellular distribution (Table S2). The C-termini ends of the pineapple SWEET proteins were less conserved and showed variable sequence lengths (Figure S1). Considering that the pineapple SWEET6, 8, and 10 are comparable to AtSWEET8, we hypothesized that they might transport glucose in a manner similar to SWEET8 of *Arabidopsis*.

### Screening of glucose transporter among AcSWEETs via homology-based search

Previous studies suggest that the conserved residues in the binding pocket of SWEET transporters play a crucial role in determining the substrate specificity of these proteins (Isoda *et al*., 2022). For example, the previously reported crystal structures of OsSWEET2b and AtSWEET13 in an inward open-facing state with their binding sites occupied by substrates provide valuable information about the transport activity of these proteins and how they interact with substrates (Yang *et al*., 2015; Wang, J. *et al*., 2022). The observed similarities between AtSWEET8 and AcSWEET10, as well as the differences with AcSWEET6 and 8, indicate that AcSWEET10 resembles more like AtSWEET8 and may exhibit a greater affinity for glucose transport compared to AcSWEET6 and 8 (Figure 2b). An analysis of the 3D structures of selected pineapple SWEET transporters compared to the AtSWEET8 transporter enables the prediction of the substrates transported by them and provides insights into their role in facilitating sugar transport within the plant (Figure 2c). Therefore, to determine that AcSWEET6, 8 and 10 would transport glucose, the 3D structures of AtSWEET8, AcSWEET6, 8, and 10 were predicted using the Alpha Fold server. The structures indicated that the proteins have a common structural arrangement with seven transmembrane segments organized into two units, connected by an inversion linker TM4. The structural arrangements of AcSWEET6, 8, and 10 were in line with previous findings in rice and *Arabidopsis* (Figure 2c). Despite the similarities, significant variations were observed in the C-terminal tails of AcSWEET6 and 8. For instance, AcSWEET8 had two additional β-sheets compared to AtSWEET8, while AcSWEET6 had an α-helix instead of β-sheets. In contrast, AcSWEET10 was structurally similar to AtSWEET8 (Figure 2c). These variations in the 3D structures of AcSWEET6 and AcSWEET8 could play a role in determining substrate specificity and transport activity and may affect the glucose transport activity of AcSWEET6 and AcSWEET8. Altogether, the results of substrate specificity and comparative structural analysis indicate that only AcSWEET10 shares a close structural similarity with AtSWEET8, suggesting that it has the highest affinity for glucose transport among AcSWEET6, 8 and 10.

### AcSWEET6, AcSWEET8, and AcSWEET10 gets localized to plasma membrane

The substrate specificity and comparative structural analysis indicated that AcSWEET6, 8 and 10 could transport glucose, so we checked their subcellular localization to confirm the putative functional location. The subcellular localizations of AcSWEET6, 8 and 10 were identified using the AcSWEET6, 8 and 10 GFP fusion proteins in tobacco epidermal cells (Figure 3). The GFP signals were compared with the earlier reported plasma membrane-localized AtSWEET8-GFP (Guan *et al*., 2008; Chen, L-Q *et al*., 2010) and the GFP control vector (35S::GFP), which was distributed throughout the cells. The fluorescence signals for AcSWEET6, 8 and 10 were observed in the plasma membrane (Figure 3). The localization results also implicated that the SWEETs show conservation for subcellular localization.

**Figure 3:**
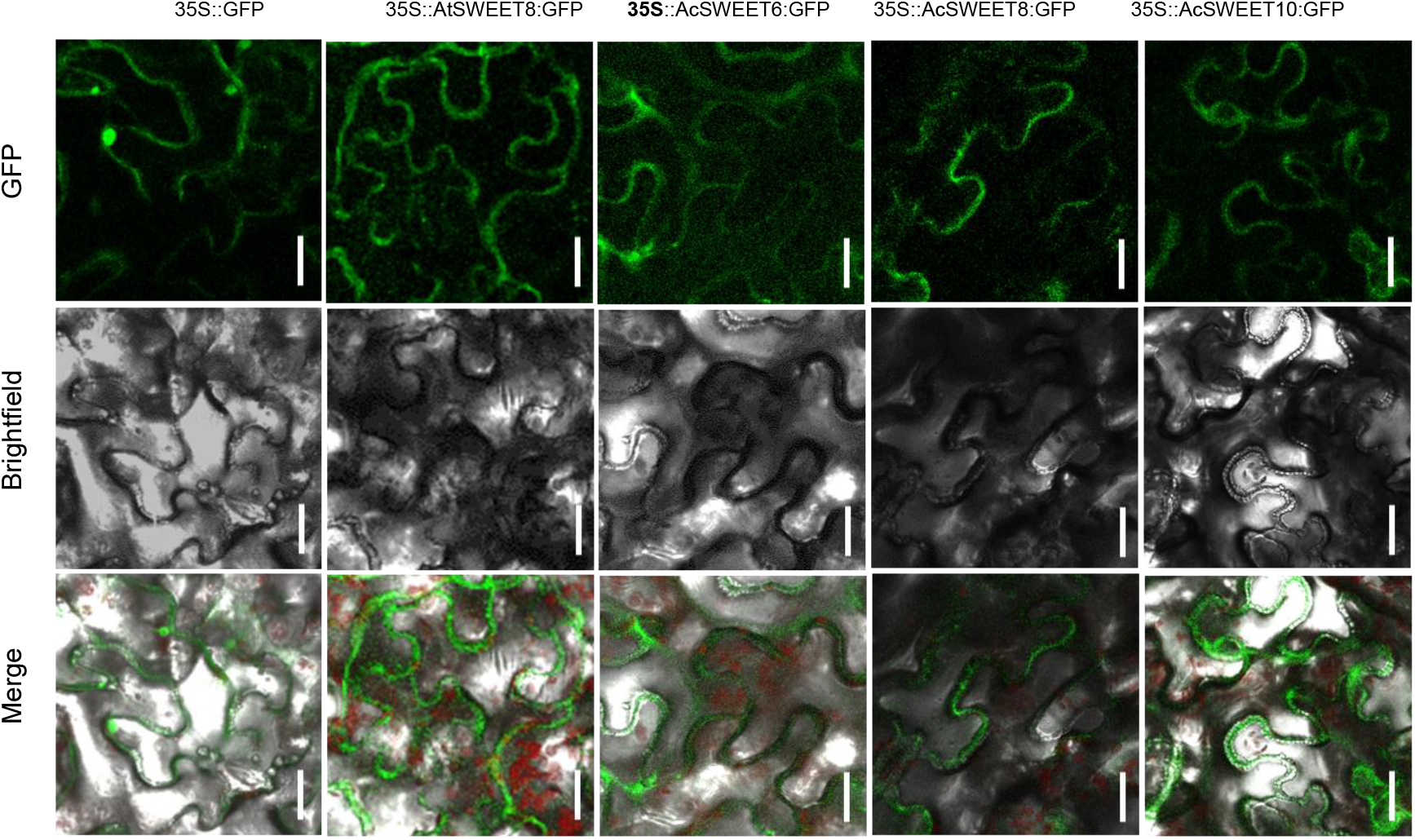
Pineapple SWEET proteins get localized to the plasma membrane. Transient subcellular localization of AcSWEET6, AcSWEET8 and AcSWEET10 proteins. The 35S::GFP, 35S::AtSWEET8:GFP, 35S::AcSWEET6:GFP, 35S::AcSWEET8:GFP and 35S::AcSWEET10:GFP fusion proteins were transiently expressed in tobacco epidermal cells and visualized by fluorescence microscopy. Scale bars: 25 µm.

### Expression of *AcSWEET* genes during pineapple pollen development

Plasma membrane specific cellular localization of AcSWEET6, 8, and 10 resembles well-characterized AtSWEET8. But, the tissue-specific localization of *AcSWEETs* still remained unknown. Therefore, we relied on the tissue-specific expression of AcSWEET8, which is observed in the stamen (Figure S2). In this context, we analyzed the expression of *AcSWEETs* using RNA-seq data from six developmental stages (S1-S6) of the stamen (Figure 4a and 4b). The results showed that 66% of the *AcSWEET* genes were upregulated during all the developmental stages of stamens. Among them, *AcSWEET6*, *AcSWEET8*, and *AcSWEET10* expressions were detected during the pineapple stamen development. The expression patterns of *AcSWEET6*, *8* and *10* were further validated using qRT PCR in the developing stamen (Figure 4c). Altogether, it suggests that pineapple SWEET transporters are similar to AtSWEET8 not only in terms of 3D structure and cellular localization but also for tissue-specific expression.

**Figure 4:**
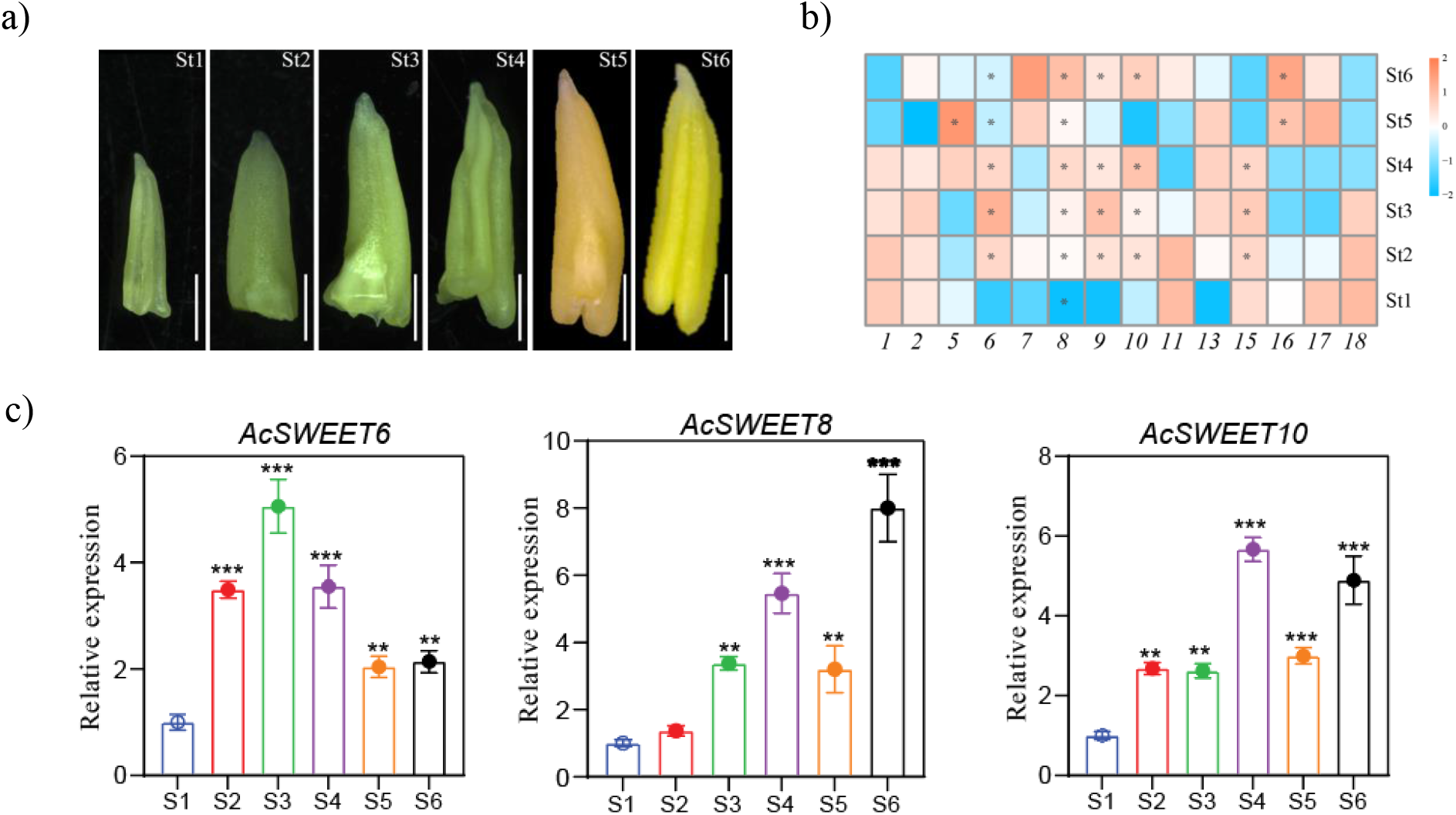
Pineapple *SWEETs* show differential expression in the anthers. (a) Morphological characteristics of the pineapple stamen tissues used for RNA-seq analysis samples were harvested at six different stages. Scale bar: 1mm. (b) Heatmap of the expression profiles of the *AcSWEETs* in stamen tissues at different stages. The heatmap was created based on the log2 (FKPM + 0.01) value of *AcSWEETs* and normalized by row. The FKPM value higher than 30 was marked with “*”. Differences in gene expression changes are shown in color as the scale. (c) Relative expression levels of *AcSWEET6,8* and *10* genes from six developmental stages of stamens. Vertical bars represent the mean ± SE of three biological replicate assays. Asterisks denote the statistical significance against Stage1 as judged by the Student’s t-test (**p < 0.01 and *** P < 0.001).

### Pineapple SWEET10 transports glucose

Although the homology-based identical pineapple SWEET transporters have similar cellular localization and tissue-specific expression, the question remained whether pineapple SWEET transporters are capable of glucose transport. Due to technical difficulties with pineapple gentic modification, it is almost impossible to knock out each glucose transporter *in plant*. As an alternative and most reliable approach, we took advantage of hexose transport-deficient yeast mutant EBY.VW4000 and performed the glucose transport assay *in vivo*. To achieve this, expression vectors harboring the coding sequences of *AcSWEET6*, *AcSWEET8*, and *AcSWEET10* were transformed into EBY.VW 4000. Transformants were allowed to grow in synthetic deficient (SD/-Trp) media supplemented with maltose (as growth control) and glucose. The yeast complementation assay showed that AcSWEET10 complements the mutant strain and grows well on glucose-supplemented media similar to AtSWEET8, while AcSWEET6 grows weakly. However, the yeast carrying AcSWEET8 and the vector (without insert) do not show detectable growth on media rich in glucose (Figure 5). These heterologous transport assays highlight that AcSWEET10 has a maximum affinity to transport glucose compared to AcSWEET6 and 8.

**Figure 5:**
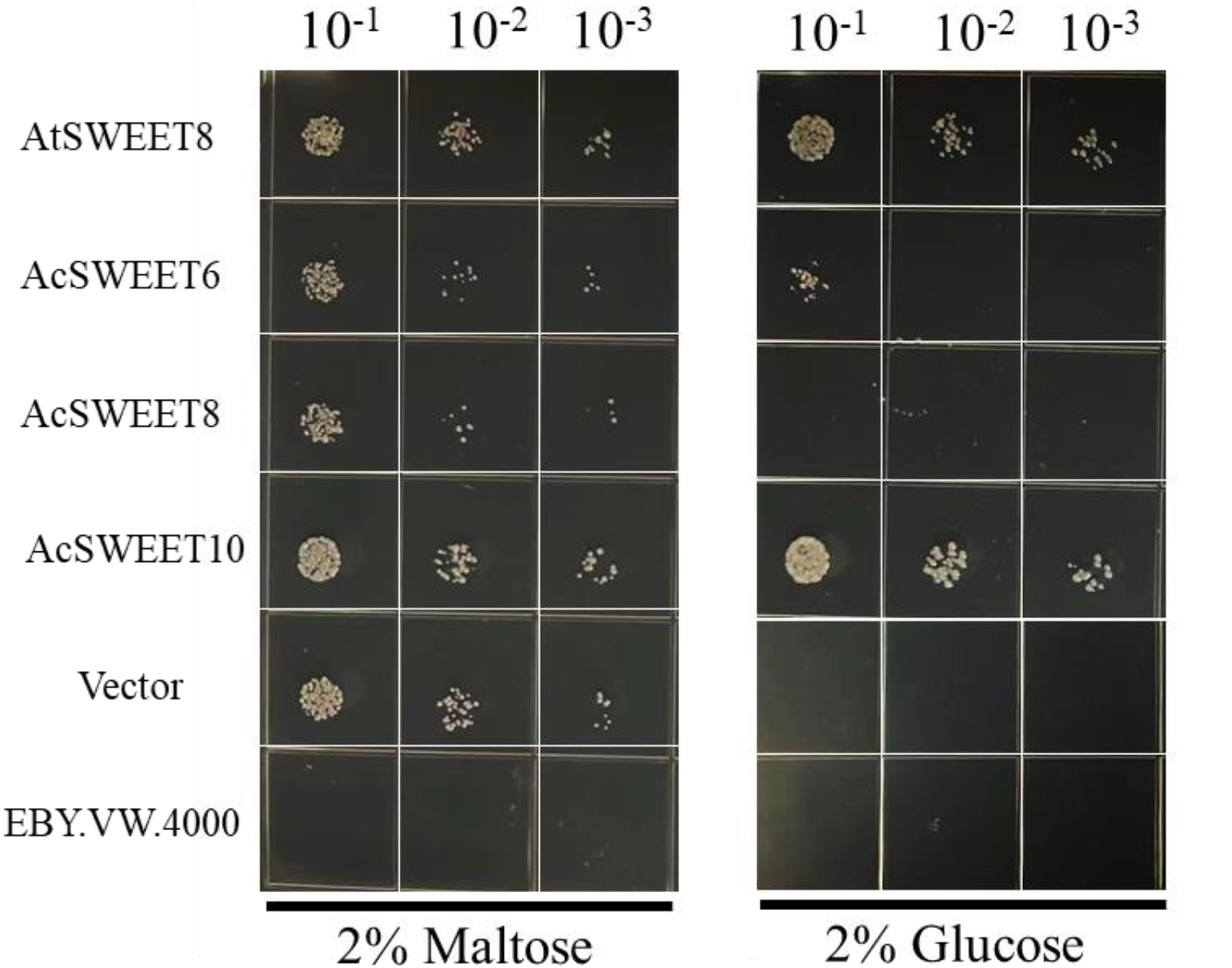
AcSWEET10 transports glucose in yeast. Glucose complementation assay of AcSWEET6, AcSWEET8 and AcSWEET10 where AtSWEET8 (positive control) and vector (without insert) (as negative control) were expressed in the hexose transport-deficient yeast mutant EBY.VW4000. The transformed yeast colonies and mutant yeast strain (as negative control) were 10-fold diluted and cultured on synthetic deficient media without tryptophan (SD/-Trp) supplemented with 2 % (w/v) maltose (as growth control) and 2 % (w/v) glucose. Images were taken after culture plates were incubated at 30°C for 4 days.

### AcSWEET10 functions as a glucose transporter in *Arabidopsis*

To investigate whether the AcSWEET10 of pineapple has a similar role in the anther development, ectopic expression of AcSWEET6, 8, and 10 in the *Atsweet8* mutant was performed to determine their complementation ability. In agreement with our substrate specificity and comparative structural analysis results, AcSWEET6 and 8 failed to complement the mutant phenotype. This was evident from pollen viability analysis using Alexander staining, which showed that most of the pollen in these lines were non-viable (Figure 6a and 6b). The percentages of pollen abortion and viable seed numbers in these lines were similar to that of the mutant (Figure 6c and 6d), indicating that AcSWEET6 and 8 do not transport glucose and therefore failed to complement the *Atsweet8* mutant. In contrast, the ectopic expression of AcSWEET10 was able to restore the defective pollen phenotype of the *Atsweet8* mutant. The AcSWEET10 expression fully rescued the low fertility phenotype of the mutant plants. The complemented lines displayed a significant improvement in silique development and seed set (Figure 7a and 7b), with the viable seed number nearly identical to that of the wild-type (Figure 7c) and a significant reduction in the pollen abortion rate in the complemented lines (Figure 7d). The anthers showed viable pollens in complemented lines, similar to the wild-type (Figure 7e). The Alexander staining showed that most of the pollens in the AcSWEET10 lines were viable, resulting in a restoration of the mutant phenotype (Figure 7f).

**Figure 6:**
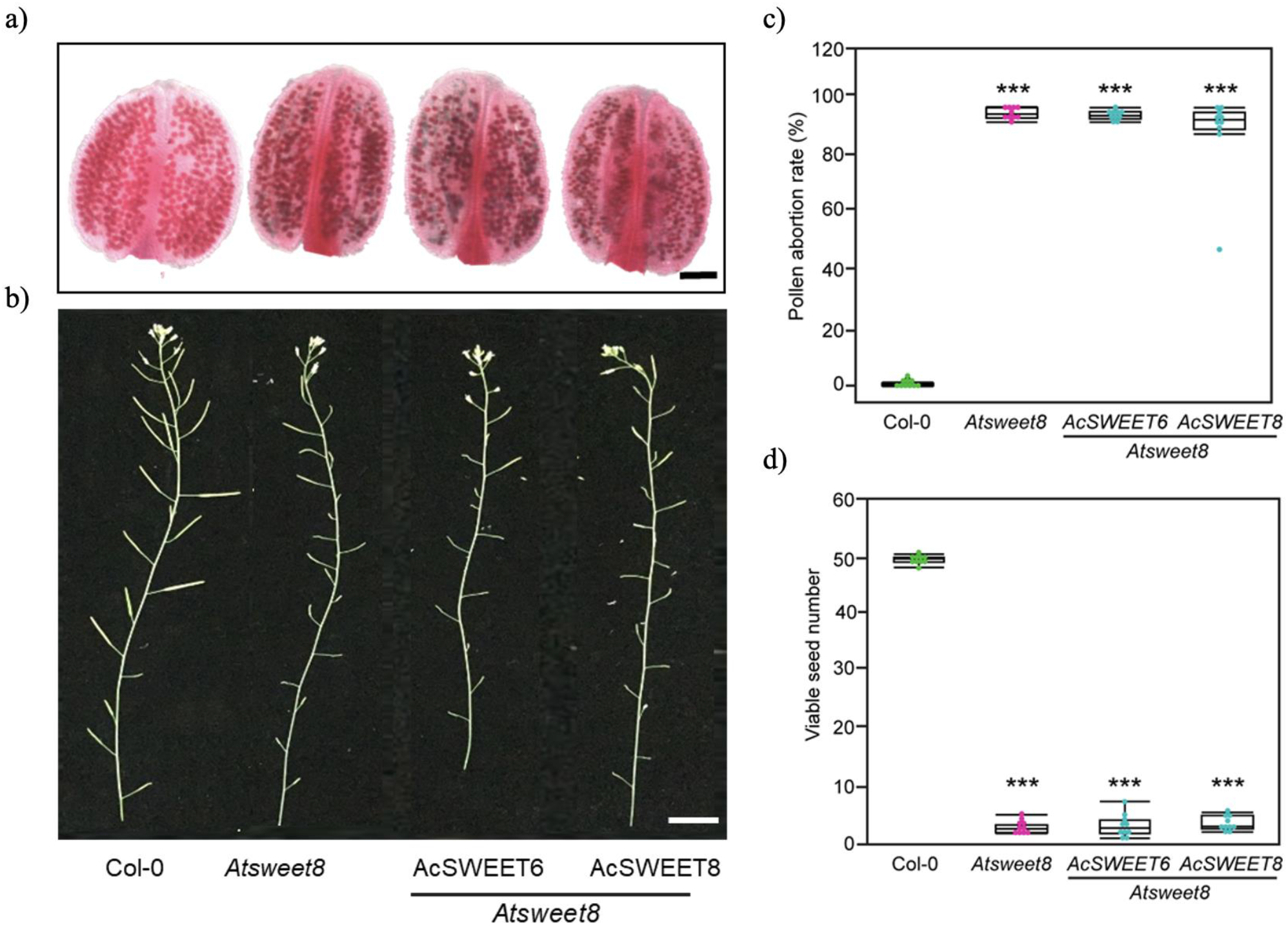
AcSWEET6 and 8 do not complement *Arabidopsis sweet8* function (a) Alexander dye-stained anthers showing non-viable pollens in non-complemented plants, similar to mutant. Scale bar: 50 µm. ( b) Primary inflorescence stems showing shorter siliques of non-complemented plants. Scale bar: 3 cm. (c) Graph showing pollen abortion rate in percentage. (d) Graph showing viable seed number. Asterisks above the columns indicate significant differences compared with Col-0.

**Figure 7:**
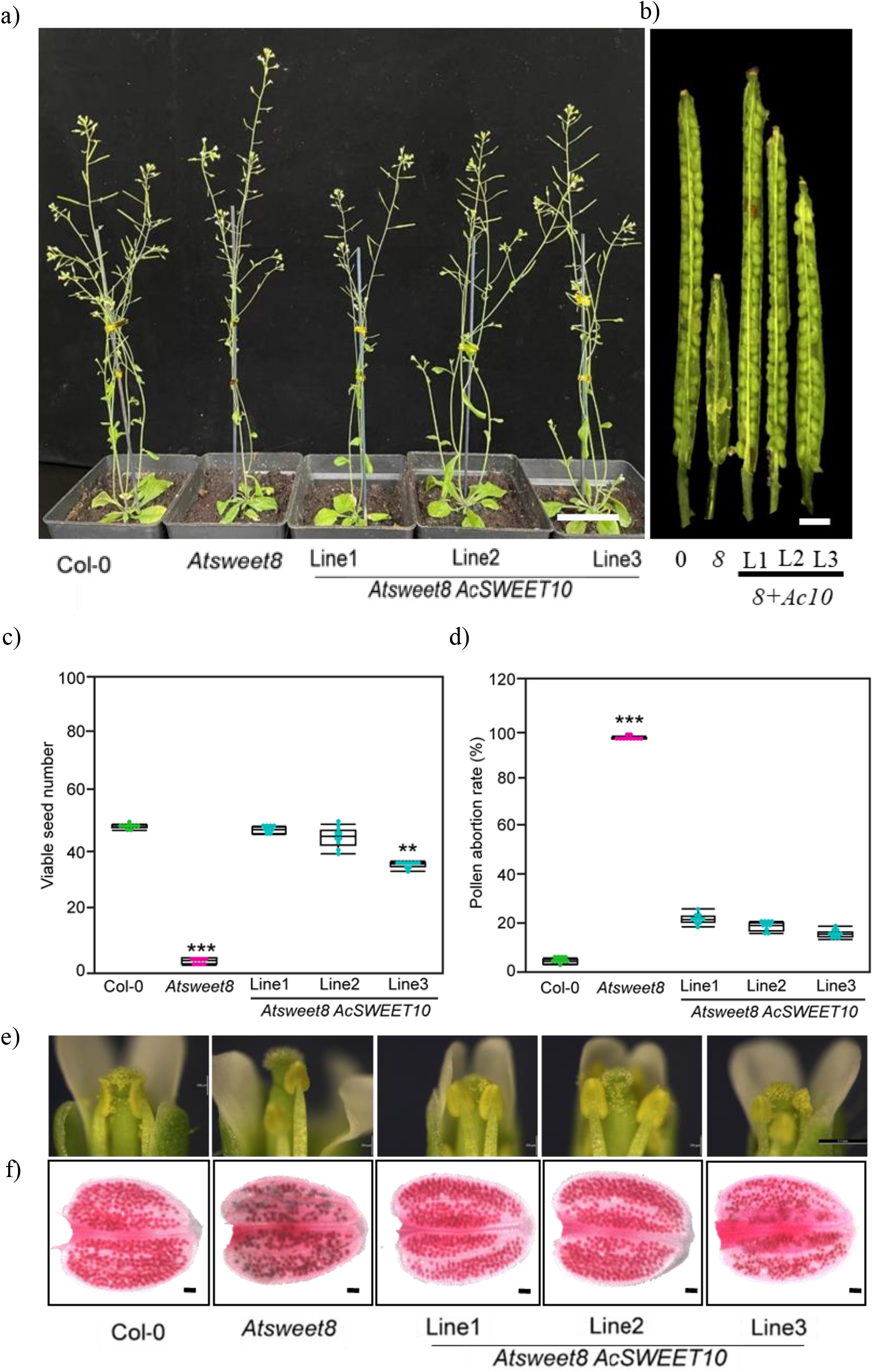
AcSWEET10 functions as a glucose transporter and complements *Arabidopsis sweet8* mutant. (a) Five-week-old plants showing silique growth from the initial stages. Scale bar: 3cm. (b) Representative photos showing variable extents of restored silique lengths in three complemented lines (L1, L2 and L3) with reference to *Atsweet8* mutant. Scale bar: 1mm. (c) Graph representing viable seed number per silique (harvested from position 5th onwards from the main inflorescence stem). (d) Graph showing pollen abortion rate in percentage. (e) Brightfield microscopy of flowers showing pollens on anthers and stigma of Col-0, *Atsweet8* and three complemented lines, the flowers showing abundant pollens on stigma of complemented lines. Scale bar: 20µm. ( f) Viable pollens in bright red stain were observed in the Alexander-stained anthers, Scale bar: 200µm). Values are represented by means + SE (n = 10 siliques per genotype) and asterisks above the columns indicate significant differences compared with Col-0.

In a previous study, the mutation of *Atsweet8* significantly reduced the expression of *CalS5* (key enzyme for callose biosynthesis), resulting in the thinning of the callose wall of the *Atsweet8* microspore (Guan *et al*., 2008; Sun *et al*., 2013). To investigate whether fertility restoration by AcSWEET10 is associated with callose deposition, we checked the deposition of callose and *CalS5* expression in the wild-type, mutant, and complemented lines. The DIC microscopy showed an improved callose wall around developing microspores at the tetrad stage in complemented lines compared to thinner degenerated walls around mutant microspores (Figure 8a). The callose fluorescence was comparable in wild-type and complemented lines in contrast to *Atsweet8* (Figure 8b and 8d). The percentage of defective tetrads has significantly reduced to up to 20% in complemented lines (Figure 8c). The relative expression of CalS5 in complemented lines was similar to that of wild-type in contrast to mutant (Figure 8e). Taken together, these results suggest that the abnormal callose deposition related to the reduced fertility in the *Atsweet8* mutant is complemented by AcSWEET10, indicating that AcSWEET10 has a similar function to SWEET8 in pineapple.

**Figure 8:**
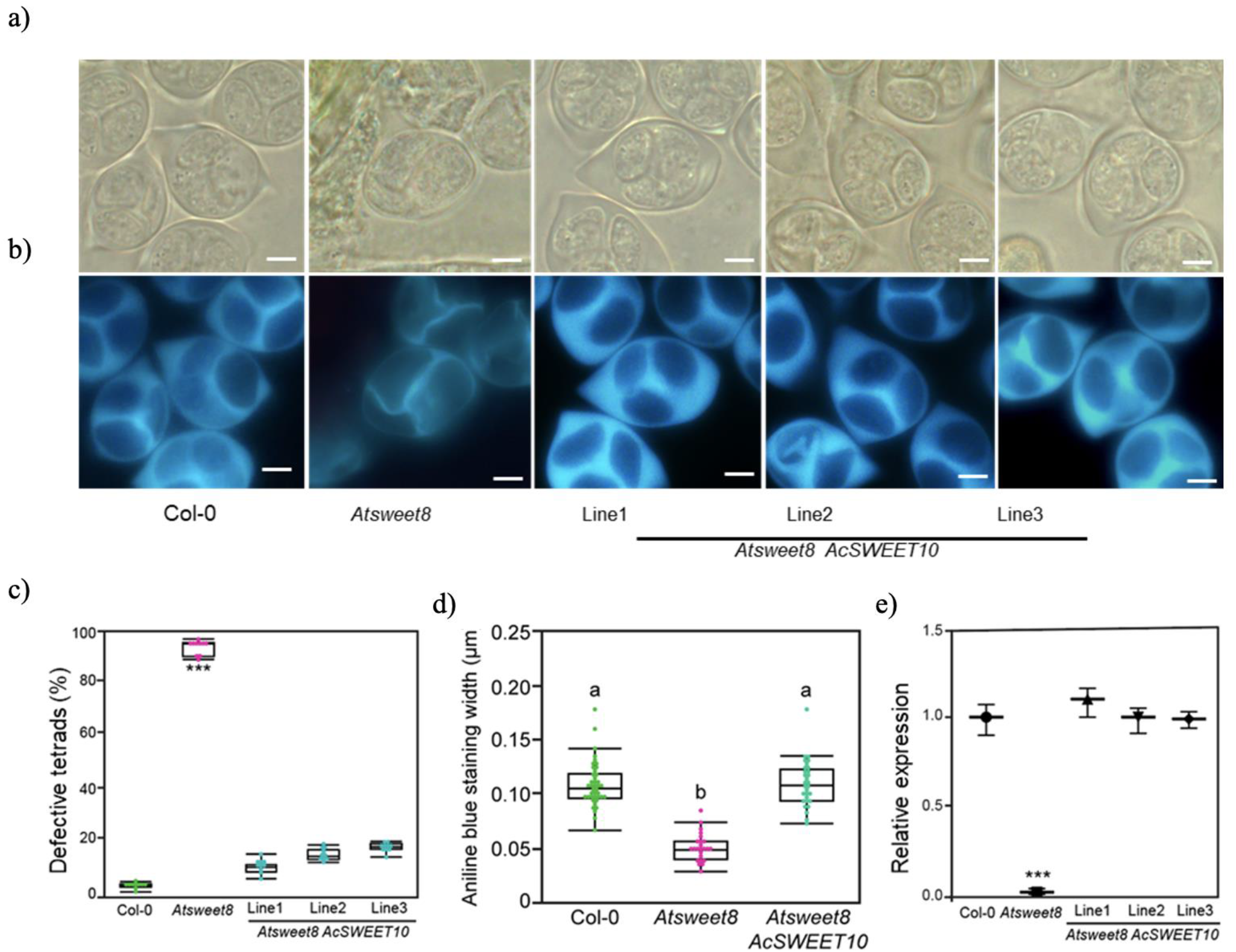
AcSWEET10 regulates callose 5 synthase levels and maintains tetrads during microspore development. (a) DIC images showing wall architecture of tetrads of wild type (Col-0), *Atsweet8* and three AcSWEET10+*Atsweet8* complemented lines. (b) Cytochemical staining with aniline blue of tetrads of in Col-0, *Atsweet8* and three complemented lines Scale bar: 20 μm. (b) Defective tetrads percentage (%) in Col-0, *Atsweet8* and three complemented lines. (d) Aniline blue staining width quantified for Col-0, *Atsweet8*, and one complemented line. The statistical test was performed based on Tukey’s Honest Test. Groups labeled with the same letter are not statistically different from each other (alpha = 0.05). (e) Relative expression levels of *callose 5 synthase* in anthers (harvested at tetrad stage) from Col-0, *Atsweet8* and three complemented lines. Values are represented by means±SE of three biological replicates. ***P<0.001.

## Discussion

The comprehensive analysis of the SWEET sugar transporter family in *Arabidopsis* and rice has provided important insights into their structure and function. In *Arabidopsis*, 17 SWEET transporters are divided into four clades, with members capable of transporting glucose or sucrose, with few exceptions (Han *et al*., 2017). In this study, 3 dimensional structures of the pineapple SWEET protein was compared with *Arabidopsis* glucose transporters to predict glucose SWEET transporters of pineapple. This prediction was further validated by exploring the glucose transport activity of AcSWEET6, 8 and 10 in yeast transport assay and functionally complementing the *Arabidopsis sweet8* mutant.

Based on sequence similarity with *Arabidopsis*, pineapple SWEETs fall into four clades, namely I, II, III, and IV. Members of these clades are potential candidates for glucose/sucrose transport. Previous studies on the structures of SWEETs have provided necessary information about their putative function in many plants (Guan *et al*., 2008; Chen, LQ *et al*., 2010; Ji *et al*., 2022). SWEETs have a single sugar-binding site within the protein and must undergo conformational changes to transport the sugar (Wang *et al*., 2014; Chen, LQ *et al*., 2015). An alignment of homologous amino acid sequences and analysis of the amino acid configuration of the binding pocket revealed a set of four residues that were conserved in subgroups with the bonafide glucose transporter (AtSWEET8) and sucrose transporter (AtSWEET13) (Figure-2a and S1). Based on these conserved residues, five sucrose transporters (AcSWEET11,12,13,15 and16) and nine glucose transporters (AcSWEET1,2, 3, 5,6,7,8,9 and10) were predicted in pineapple (Figure 2a).

*Arabidopsis* SWEET proteins play critical roles in the transport of glucose directly or in the form of sucrose between compartments, cells, and organs (Chen, LQ *et al*., 2010; Sun *et al*., 2013; Eom *et al*., 2015). AtSWEET8 is a well-known plasma membrane glucose transporter, and its loss of function leads to severely defective pollen and a reduction of up to 90% in the seed set (Guan *et al*., 2008); therefore, it was utilized as the reference for this study. Clade III members (AcSWEET6, 8 and 10) displayed high similarity in their physicochemical properties with the AtSWEET8 and localize to the plasma membrane (Figure 3). The glucose molecule follows a specific path by interacting with residues across the SWEET8 cavity (Selvam *et al*., 2019). The same residues for glucose entry and exit in AcSWEET 6, 8 and 10 structures indicated their potential for glucose transport activity (Figure S1). Besides, similar to *AtSWEET8*, the high expression of *AcSWEET6*, *8* and *10* in pineapple stamens showed their potential role in sugar supply in male reproductive organs (Figure 4a, 4c and S2).

The 3D structure of AcSWEET6, 8, and 10 proteins using the AlphaFold algorithm showed high similarity in their overall structure, but they significantly differed at the C-terminal region (Figure 2c). The predicted structures suggest that the C-terminal region of AcSWEET6 forms an extra alpha-helix, which is absent in AcSWEET8 and 10. In addition, the C-terminal region of AcSWEET8 possessed an additional β-sheet which changed the structure leading to the hindrance of the pore (Figure 2c). Previously, it was shown that three mutant alleles of the *Atsweet8* gene (Salk_142803, Salk_062567and Salk_092239) with a T-DNA insertion in the first exon, fifth exon and first intron, respectively (Figure S3). However, the stronger defect in phenotype pollen of the *AtSWEET8* gene was observed in the *rpg1* mutant, a T-DNA insertion in the last intron. Consequently, in RT-PCR of the *Atsweet8* mutant, the T-DNA insertion primarily affected the transcription of the last exon, which corresponds to the C-terminal tail of its protein (Guan *et al*., 2008). Altogether this suggests that the C-terminal region of the SWEET8 plays an essential role in its function. Besides, several studies indicate the significance of C-terminal regions for the functions of different transporters (Nobukuni *et al*., 2009; Yamada *et al*., 2017; Fujita *et al*., 2018). The structural difference of the AcSWEET6 and 8 at the C-terminal compared to AcSWEET10 and AtSWEET8 could impact the glucose transport activity of the proteins AcSWEET6 and 8. Consistently, the present study revealed that AcSWEET10 has a structure similar to AtSWEET8 and displayed higher glucose transport activity than SWEET6 and 8 (Figure 2b, 2c and 5), most likely due to the difference in their C-terminal region, which may be hindering the sugar transport.

Experimental verification of the putative transporters using multiple hexose transporter-deficient yeast showed that only AcSWEET10 could enable robust growth on glucose (Figure 5). Interestingly, AcSWEET10 retained a conserved structure for glucose transport and demonstrated complementation of the *Atsweet8* mutant (Figure 2b, 2c and 7). Although AcSWEET6 and 8 have identical conserved residues, their failure to transport glucose and complement the mutation, indicates the importance of the C-terminal for their function (Figure 6).

To summarize, this research has highlighted the significance of the C-terminal region for the appropriate functioning of SWEETs. The results of the study revealed that AcSWEET10 has a conserved function and imitates the overall structure of AtSWEET8, implying that residues in the C-terminal region can have an impact on transporter activity (Figure 9). Further investigation is needed to explore the C-terminal regions of SWEET transporters and the possible presence of motifs interacting with glucose, particularly in the case of AcSWEET6 and 8. Overall, these findings have important implications for understanding plant physiology and metabolism and developing strategies to improve crop yield and quality. Further investigation in this area could develop novel tools for manipulating sugar transport in plants, which could have far-reaching benefits for agriculture and food security.

**Figure 9:**
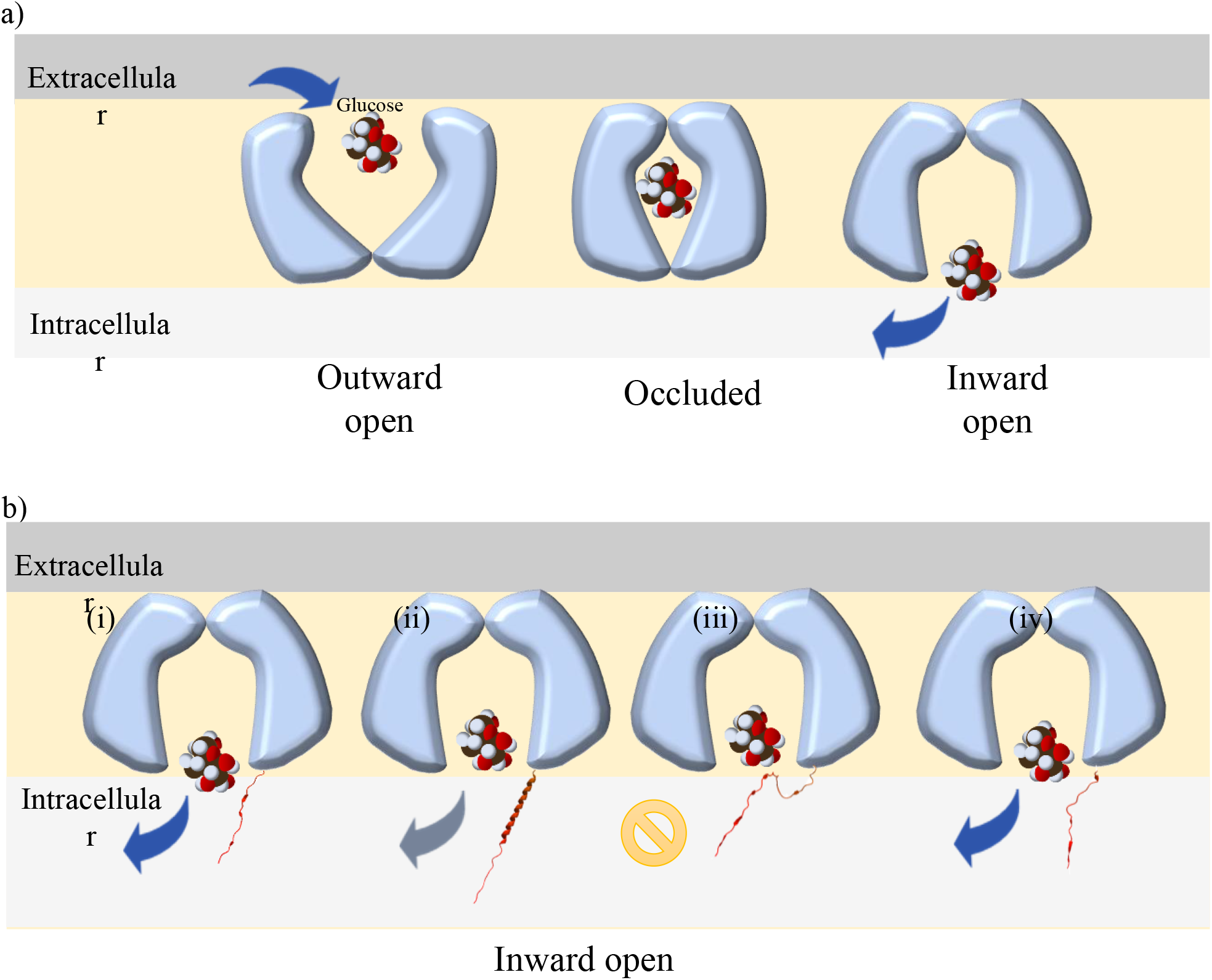
A schematic model representing the glucose transport mechanism in SWEET8 proteins. (a) model representing three different conformations of the SWEET transporter (outward open, occluded and inward open) showing glucose transport according to Selvam, et al. 2019, (b) model representing inward open conformations of AtSWEET8 (i), AcSWEET6 (ii), AcSWEET8 (iii), and AcSWEET10 (iv) proteins with arrows (blue: strong, grey:weak and stop signal: restricted) representing exit of glucose molecule through the transporter.

## Acknowledgments

We especially thank Dr. Binghua Wu (Fujian Agriculture and Forestry University, China) for kindly providing the yeast mutant strain EBY.VW4000 and Prof. Zhong-Nan Yang (Shanghai Normal University, China) for sharing *Atsweet8* seeds.

## Author contributions

M.A. and Y. Q. designed the study. B.F., L.W., X.W. and P.Z performed the experiments. B.F, M.A.A., M.A., and Y.Q. analyzed the data and wrote the paper.

## Data availability

The data and materials supporting this study are available upon request from the corresponding author.

## Supplemental Figure and Table

**Table S1:** Pairwise comparison of the amino acid identity of Pineapple SWEET (18), AtSWEET13, AtSWEET8 and OsSWEET2b.

**Table S2:** Physicochemical properties of pineapple SWEETs and AtSWEET8.

**Table S3:** List of the primers used in the study.

**Figure S1:** Multiple sequence alignments of SWEET proteins of pineapple. AtSWEET8 and AtSWEET13 serve as the reference. The numbers on top of the table represent AtSWEET13.

**Figure S2:** Histochemical GUS localization of AtSWEET8 in which GUS expression was driven with the SWEET8 promoter with the *SWEET8* gene. Three independent lines were analyzed with similar results. Scale bar: 1 mm

**Figure S3:** Schematic representation of T-DNA insertion sites in the *SWEET8* gene of *Arabidopsis*.

