## Additional files for "Substrate specificity and functional conservation of SWEET10 transporter in pineapple"

**Table S1:** Pairwise comparison of the amino acid identity of Pineapple SWEET (18), AtSWEET13, AtSWEET8 and OsSWEET2b.

| | AcSWEET4 | AtSWEET8 | AcSWEET9 | AcSWEET7 | AcSWEET5 | AcSWEET6 | AcSWEET8 | AcSWEET10 | AcSWEET11 | AtSWEET13 | AcSWEET13 | AcSWEET14 | AcSWEET12 | AcSWEET15 | AcSWEET16 | AcSWEET3 | AcSWEET17 | AcSWEET18 | AcSWEET2 | AcSWEET1 | AcSWEET2b |
|---|---|---|---|---|---|---|---|---|---|---|---|---|---|---|---|---|---|---|---|---|---|
| 1: AcSWEET4 | | 27.59 | 23.42 | 26.09 | 27.10 | 28.28 | 32.63 | 31.82 | 23.14 | 18.11 | 23.01 | 27.18 | 25.49 | 23.64 | 25.41 | 25.74 | 25.19 | 10.34 | 32.35 | 26.61 | 26.14 |
| 2: AtSWEET8 | | | 33.48 | 38.46 | 39.57 | 44.78 | 44.35 | 43.67 | 34.84 | 33.04 | 33.48 | 20.69 | 32.58 | 32.00 | 33.48 | 34.06 | 39.64 | 35.33 | 30.99 | 39.37 | 30.63 |
| 3: AcSWEET9 | | | | 32.95 | 34.32 | 37.25 | 54.80 | 45.30 | 26.64 | 25.70 | 27.27 | 19.12 | 26.67 | 26.09 | 26.90 | 28.80 | 29.62 | 28.77 | 31.98 | 32.53 | 34.55 |
| 4: AcSWEET7 | | | | | 40.16 | 45.81 | 67.12 | 57.77 | 29.54 | 25.31 | 25.33 | 18.14 | 27.95 | 25.44 | 25.00 | 25.55 | 30.65 | 36.81 | 24.84 | 33.04 | 27.27 |
| 5: AcSWEET5 | | | | | | 60.16 | 47.81 | 55.32 | 29.60 | 26.40 | 27.16 | 19.74 | 29.79 | 27.89 | 28.74 | 29.96 | 32.39 | 32.43 | 31.11 | 35.08 | 32.89 |
| 6: AcSWEET6 | | | | | | | 54.98 | 60.00 | 32.47 | 31.65 | 32.30 | 24.40 | 36.04 | 31.90 | 35.96 | 30.33 | 38.03 | 36.11 | 33.14 | 40.95 | 37.44 |
| 7: AcSWEET8 | | | | | | | | 68.09 | 34.36 | 34.33 | 32.44 | 24.76 | 34.23 | 32.75 | 35.11 | 32.08 | 41.48 | 37.50 | 31.95 | 43.17 | 36.53 |
| 8: AcSWEET10 | | | | | | | | | 36.74 | 34.39 | 33.02 | 27.04 | 34.88 | 34.25 | 37.67 | 31.58 | 43.78 | 38.19 | 34.52 | 43.78 | 37.61 |
| 9: AcSWEET11 | | | | | | | | | | 47.33 | 51.38 | 29.17 | 52.89 | 50.00 | 48.65 | 31.84 | 36.78 | 36.91 | 31.21 | 33.20 | 31.25 |
| 10: AtSWEET13 | | | | | | | | | | | 52.59 | 30.30 | 51.89 | 54.05 | 51.06 | 30.92 | 35.94 | 36.00 | 30.46 | 36.14 | 34.22 |
| 11: AcSWEET13 | | | | | | | | | | | | 34.65 | 56.86 | 58.75 | 58.39 | 33.19 | 32.20 | 33.56 | 29.76 | 31.67 | 31.05 |
| 12: AcSWEET14 | | | | | | | | | | | | | 43.68 | 43.57 | 65.25 | 23.42 | 24.90 | 14.48 | 23.87 | 21.49 | 20.30 |
| 13: AcSWEET12 | | | | | | | | | | | | | | 72.06 | 63.70 | 36.21 | 38.22 | 38.26 | 32.35 | 37.13 | 31.67 |
| 14: AcSWEET15 | | | | | | | | | | | | | | | 71.65 | 34.55 | 37.40 | 35.57 | 32.76 | 36.18 | 30.67 |
| 15: AcSWEET16 | | | | | | | | | | | | | | | | 30.99 | 35.36 | 32.89 | 34.12 | 36.03 | 34.39 |
| 16: AcSWEET3 | | | | | | | | | | | | | | | | | 34.43 | 32.67 | 38.71 | 39.75 | 37.99 |
| 17: AcSWEET17 | | | | | | | | | | | | | | | | | | 52.67 | 34.71 | 43.03 | 36.49 |
| 18: AcSWEET18 | | | | | | | | | | | | | | | | | | | 31.25 | 39.19 | 33.11 |
| 19: AcSWEET2 | | | | | | | | | | | | | | | | | | | | 44.25 | 50.00 |
| 20: AcSWEET1 | | | | | | | | | | | | | | | | | | | | | 45.18 |
| 21: OsSWEET2b | | | | | | | | | | | | | | | | | | | | | |

**Table S2:** Physicochemical properties of pineapple SWEETs and AtSWEET8.

| Gene name | Gene ID | Length (aa) | MW(Da) | pI | I.I. | A.I. | GRAVY | Sub-loc. | TMH |
|---|---|---|---|---|---|---|---|---|---|
| AcSWEET1 | Aco011302 | 258 | 28.902 | 9.21 | 34.93 | 107.64 | 0.519 | PM | 7 |
| AcSWEET2 | Aco016508 | 194 | 22.292 | 7.67 | 32.41 | 114.48 | 0.664 | PM | 3 |
| AcSWEET3 | Aco010708 | 288 | 32.685 | 9.34 | 46.28 | 106.98 | 0.45 | PM | 7 |
| AcSWEET4 | Aco006346 | 175 | 20.285 | 8.47 | 42.09 | 75.77 | -0.393 | Chlo | 1 |
| AcSWEET5 | Aco005793 | 288 | 32.614 | 9.75 | 44.63 | 113.68 | 0.445 | PM | 6 |
| AcSWEET6 | Aco004463 | 256 | 28.143 | 9.16 | 39.03 | 113.75 | 0.654 | PM | 6 |
| AcSWEET7 | Aco006158 | 885 | 98.972 | 9.47 | 45.09 | 96.84 | 0.006 | PM | 8 |
| AcSWEET8 | Aco006156 | 251 | 27.396 | 9.2 | 42.77 | 124.98 | 0.885 | PM | 7 |
| AcSWEET9 | Aco006155 | 423 | 45.779 | 5.29 | 71.79 | 95.04 | 0.192 | PM | 7 |
| AcSWEET10 | Aco016418 | 235 | 25.985 | 8.58 | 41.55 | 132.3 | 0.927 | PM | 7 |
| AcSWEET11 | Aco001900 | 268 | 29.775 | 9.01 | 37.7 | 121.12 | 0.703 | PM | 7 |
| AcSWEET12 | Aco019048 | 281 | 31.573 | 8.93 | 40.49 | 121 | 0.577 | PM | 7 |
| AcSWEET13 | Aco004628 | 274 | 30.92 | 7.66 | 43.75 | 124.45 | 0.777 | PM | 7 |
| AcSWEET14 | Aco016039 | 283 | 31.731 | 9.19 | 56.81 | 94.38 | -0.068 | V | 3 |
| AcSWEET15 | Aco003627 | 265 | 29.65 | 9.71 | 27.51 | 127.25 | 0.773 | PM | 7 |
| AcSWEET16 | Aco017831 | 302 | 33.36 | 6.53 | 40.34 | 118.34 | 0.549 | PM | 7 |
| AcSWEET17 | Aco002476 | 293 | 32.113 | 9.5 | 31.97 | 116.08 | 0.499 | V | 7 |
| AcSWEET18 | Aco006347 | 150 | 16.29 | 9.4 | 32.62 | 143.53 | 1.155 | V | 4 |
| AtSWEET8 | At5G40260 | 240 | 26.8 | 9.6 | 41 | 121.38 | 0.764 | PM | 7 |

PM- plasma membrane; Chlo- chloroplasts; V-Vacuole

*Legends: aa -amino acid; MW- molecular weight in daltons (Da); pI- isoelectric point; I.I. -instability index; GRAVY- grand average of hydropathy; Loc- subcellular localization; TMH- transmembrane helices.

**Table S3:** List of the primers used in the study.

| Name | Primer Sequence (5'---> 3') | |
|---|---|---|
| | Forward | Reverse |
| **A. RT-qPCR** | | |
| ***AtSWEET8*** | GTGGTGGCTATCATTCTTAT | TCCAGCATTAACGAAACAGA |
| ***AcSWEET5*** | CCTATGCTCTCATCCGCTTC | CCTCTGCCAGACTCAACTCC |
| ***AcSWEET6*** | CACCAGCGCCAACTTTTTAT | ACGAGGAAGAGGAGCACGTA |
| ***AcSWEET7*** | AGCAGCAGCAGCAGTTACAA | CATGCTTCAATTGCTCCTCA |
| ***AcSWEET8*** | GGTGATTACTGGGGTGATGG | CGGATGAAGGCATAGATCGT |
| ***AcSWEET9*** | CCTACCTCGTGACGCTCTTC | ACGACAACTCCGCAGCTACT |
| ***AcSWEET10*** | CTGGACCACCTATGCCCTAA | CAAGAGTCTTCGAGGGCAAC |
| ***AcEF1a*** | TCTTCTCAGGGAAGGTCTCTAC | CTCTGCACACTCTTCACATACA |
| **B. Cloning** | | |
| ***AtSWEET8*** | CACCATGGTTGATGCAAAACAAGT | AACCCTCTCCGTAGCAGAAAT |
| ***AcSWEET6*** | CACCATGGTTTCTGCTGATACCATC | GGTTTGAGCTTTTTGGCC |
| ***AcSWEET8*** | CACCATGGTCTCCGCCGACGCAGTT | TTTATCTACTGTGATAGTGGC |
| ***AcSWEET10*** | CACCATGACTGATCCTGATACTATCC | GACTTCAAGAGTCTTCGAGG |
| **C. Yeast complementation assay** | | |
| ***AtSWEET8*** | ATGGTTGATGCAAAACAAGTTCGT | AACCCTCTCCGTAGCAGAAAT |
| ***AcSWEET6*** | ATGGTTTCTGCTGATACCATC | GGTTTGAGCTTTTTGGCC |
| ***AcSWEET8*** | ATGGTCTCCGCCGACGCAGTT | TTTATCTACTGTGATAGTGGC |
| ***AcSWEET10*** | ATGACTGATCCTGATACTATCC | GACTTCAAGAGTCTTCGAGG |

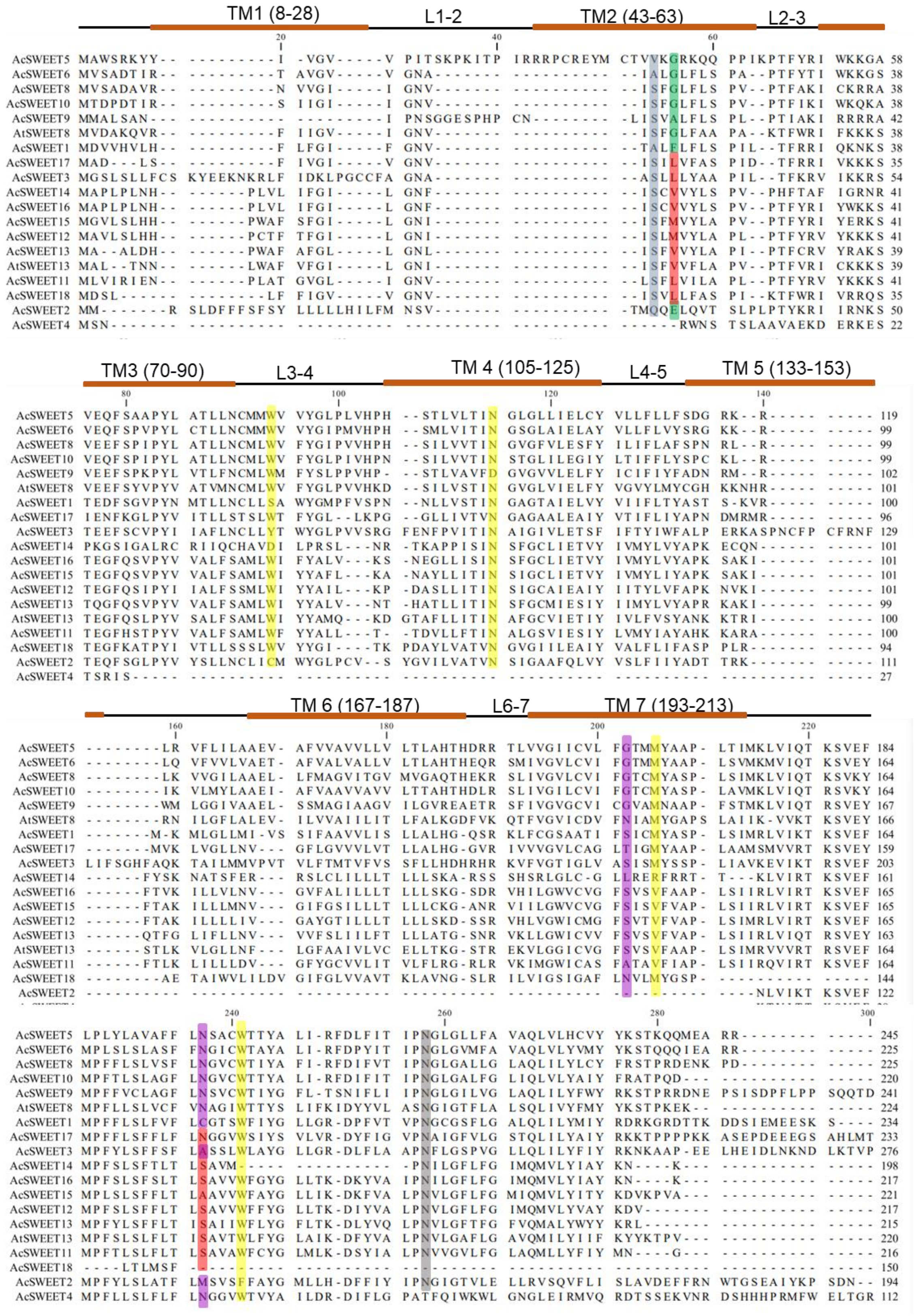


**Figure S1:** Multiple sequence alignments of SWEET proteins of pineapple. AtSWEET8 and AtSWEET13 serve as the reference. The numbers on top of the table represent AtSWEET13.

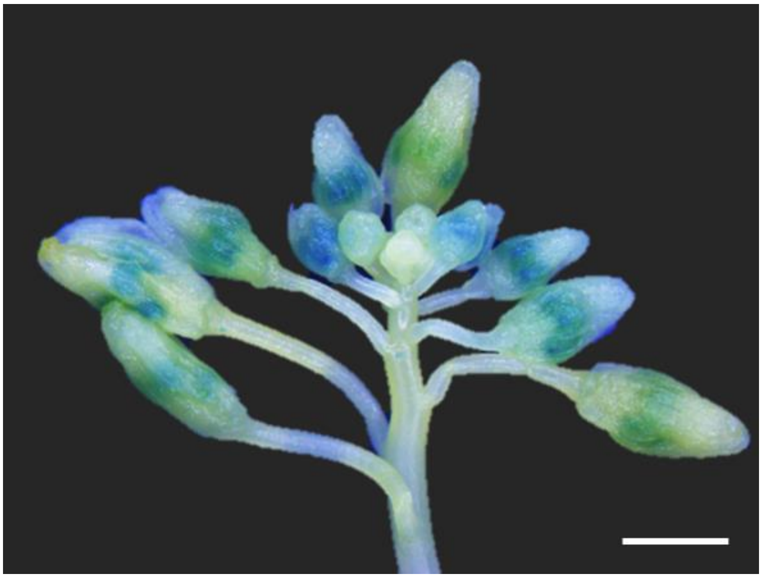

**Figure S2:** Histochemical GUS localization of AtSWEET8 in which GUS expression was driven with the SWEET8 promoter with the *SWEET8* gene. Three independent lines were analyzed with similar results. Scale bar: 1 mm

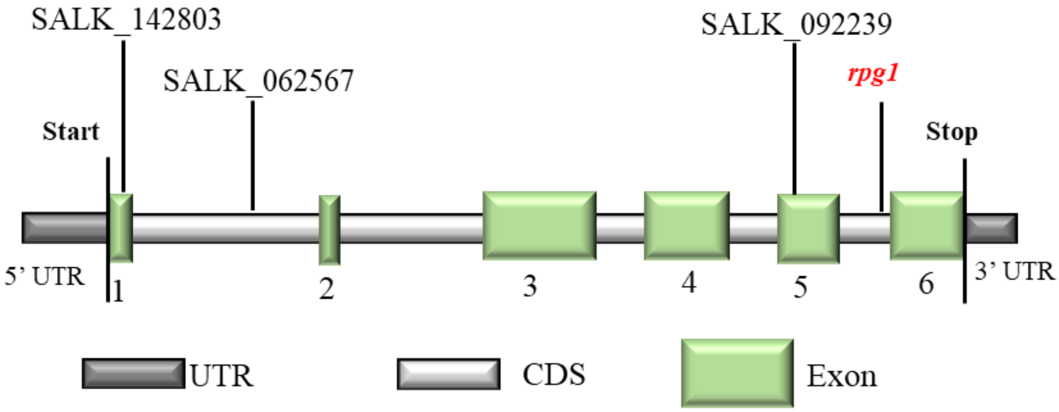


**Figure S3:** Schematic representation of T-DNA insertion sites in the *SWEET8* gene of *Arabidopsis*.
